# Circuit firing homeostasis following synaptic perturbation ensures robust behavior

**DOI:** 10.1101/2024.08.27.609984

**Authors:** Adam Hoagland, Zachary Louis Newman, Zerong Cai, Ehud Y. Isacoff

## Abstract

Homeostatic regulation of excitability and synaptic transmission ensures stable neural circuit output under changing conditions. We find that pre- or postsynaptic weakening of motor neuron (MN) to muscle glutamatergic transmission in *Drosophila* larva has little impact on locomotion, suggesting non-synaptic compensatory mechanisms. *In vivo* imaging of MN to muscle synaptic transmission and MN activity both show that synaptic weakening *increases* activity in tonic type Ib MNs, but not in the phasic type Is MN that innervate the same muscles. Additionally, an inhibitory class of pre-MNs that innervates type Ib—but not Is—MNs *decreases* activity. Our experiments suggest that weakening of MN evoked synaptic release onto the muscle is compensated for by an increase in MN firing due to a combined cell-autonomous increase in excitability and decreased inhibitory central drive. Selectivity for type Ib MNs may serve to restore tonic drive while absence of firing adjustment in the convergent Is MN can maintain the contraction wave dynamics needed for locomotion.

## Introduction

A fundamental property of the nervous system is its ability to support behavior across a large set of changing conditions. This robustness emerges from mechanisms that stabilize circuit function as connections are formed and eliminated, change synaptic strength due to activity-dependent plasticity, and adjust to physiological or pathological perturbations (Marder, 2006). Such compensatory responses ensure reliable circuit output by adjusting both pre and postsynaptic function (Marder, 2007). Postsynaptic homeostatic mechanisms can rebalance circuits by either adjusting postsynaptic strength via a scaling mechanism that preserves the relative strength of synapses, or by adjusting postsynaptic cell excitability (Turrigiano, 2012; Wen & Turrigiano, 2024). In the glutamatergic type I motor neuron to muscle synapse of the larval *Drosophila* neuromuscular junction (NMJ), weakening of the postsynaptic response (by mutation of the GluRIIA subunit of the postsynaptic ionotropic glutamate receptor or its partial pharmacological block) triggers presynaptic homeostatic plasticity (PHP), a compensatory boost in transmitter release that is triggered by retrograde signaling (Peterson, 1997; Frank, 2006; Davis, 2015; Nair, 2021).

*Drosophila* larvae have two classes of type I glutamatergic motor neurons (MNs) that convergently innervate each muscle: a tonic, ramp-bursting type Ib MN and a phasic, abrupt-bursting type Is MN, which provide approximately half of the excitatory drive to each muscle (Lee et al., 2009). The type Ib MN partially compensates for weakened transmission, while the type Is MN input does not compensate through changes in action potential (AP)-evoked presynaptic release (Newman, 2017; Genç and Davis, 2019). This apparent shortfall in compensation suggests that there should be a defect in circuit output and locomotion. However, we find here that locomotion in GluRIIA mutant larvae is close to normal. To understand how locomotion could be preserved when synaptic transmission per AP remains reduced, we examined locomotion in larvae with severely reduced glutamate release due to knockdown of each of three key components of the MN transmitter release machinery: the MN voltage-gated Ca^2+^ channel, cac, which couples the AP to release, the vesicle priming protein unc-13, and RIM binding protein (Rbp). Each knockdown greatly reduced the probability of AP-evoked release (*P_r_*), leading us to expect a major disruption in locomotion. However, once again, locomotion was near normal. This suggested that a second form of homeostatic adjustment makes up the difference when PHP is insufficient to fully compensate for reduced transmitter release. We examined MN activity (using cytoplasmic GCaMP6f) and glutamatergic synaptic transmission (using SynapGCaMP6f, our postsynaptic density targeted GCaMP6f that measures Ca^2+^ influx through the GluRII receptors) in larvae undergoing restrained locomotion (Newman et al, 2017). We find that animals with presynaptically or postsynaptically weakened neuro-muscular transmission *increase* the duration of the bouts of MN activity and synaptic transmission that drive locomotory peristaltic waves of muscle contraction, but only in type Ib MNs. This suggests a form of compensation for synaptic weakening that is accomplished by an increase in either type Ib MN excitability or in central drive specifically to type Ib MNs.

In addition, Ca^2+^ imaging with cytoplasmic GCaMP6m in central pre-motor neurons (PMNs) that innervate type Ib (but not type Is) MNs revealed a second effect of disrupted MN-muscle transmission: *decreased* activity in the period-positive median segmental interneurons (PMSIs). Thus, weakening of per AP glutamatergic transmission appears to be compensated by increased AP firing of type Ib MNs due to a change in the activity of a central input to type Ib MNs. Lack of synaptic compensation via PHP or of firing compensation in the type Is input suggests that type Ib MNs are likely setting and maintaining the tone of transmission allowing type Is MNs to remain in control the timing of muscle contraction in order to ensure orderly locomotory patterns.

## Results

### Screen to identify presynaptic deficiency in neurotransmission at Ib and Is synapses

In PHP, reduced current through postsynaptic GluRIIs (reduction in quantal size), due to either a genetic perturbation (the null mutation of the GluRIIA subunit, GluRIIA^-/-^) or pharmacological block of the GluRII pore with the wasp venom philanthotoxin, is compensated for by an increase in the total amount of glutamate released per AP (increase in quantal content or number of vesicles released) (Davis and Müller, 2015). Because PHP occurs at Ib synapses but not Is synapses and is incomplete even at Ib synapses (Newman, 2017; Genç and Davis, 2019), the compensation is only partial. This suggests that an additional homeostatic mechanism may be needed to ensure locomotor output. In search of such a mechanism, we first explored other synaptic perturbations that weaken MN to muscle synaptic transmission by turning from postsynaptic to a direct presynaptic perturbation. We evaluated the effects of these disruptions with optical quantal analysis to measure synaptic transmission at hundreds individual synapses simultaneously and from both Ib and Is inputs.

To perturb presynaptic transmitter release, we utilized a MN-specific Gal4 driver, OK6 (Aberle 2002), to conditionally drive expression in Ib and Is MNs of RNAis targeting genes that encode proteins that are part of the transmitter release machinery. While the selected proteins are known to impact total quantal content (as measured electrophysiological for total release), their impact on transmission specially at Ib versus Is synapses has not been resolved. We tested: (i) Brp, the ELKS family AZ scaffolding protein, (ii) Cacophany (cac), the voltage-gated Ca^2+^ channel that couples the AP to transmitter release, (iii) Synaptotagmin 1 (Syt1), the Ca^2+^ sensor for AP-evoked release, (iv) Rab3, the vesicular small GTPase, (v) the Rab3 interacting molecule (RIM), which clusters Ca^2+^ channel at the release site, (vi) the RIM-binding protein (Rbp) that connects release sites to Brp, (vii and viii) the regulators of synaptic vesicle priming unc-13 and unc-18, (ix) the synaptic vesicle glutamate transporter (VGluT), (x) the AZ scaffolding protein Liprin-*αα*, (xi) leukocyte antigen-related (LAR), a receptor tyrosine phosphatase, and (xii) the Shaker potassium channel (Sh) (Cunningham 2023; Rizalar 2021; Südhof TC 2012; Xie 2012).

Synaptic transmission was first imaged in the dissected preparation, where the brain and ventral nerve cord (VNC) are removed and nerve bundles containing Ib and Is MN axons are stimulated electrically. We visualized AP-evoked inward postsynaptic currents by detecting the Ca^2+^ rise due to influx through the GluRII postsynaptic receptors with SynapGCaMP6f, a genetically-encoded reporter that localizes GCaMP6f to the postsynaptic density **(Fig. 1A**) (Guerrero, 2005; Reiff, Ball, Peled, 2011, 2014). We identified synapses from spatial clusters of AP-evoked optical quantal events **(Fig. 1B**) (Newman, 2017). We measured quantal density, the number of evoked transmission events per stimulus per unit area of postsynaptic membrane (the optical equivalent of quantal content). As observed earlier (Newman, 2017; 2022), in control animals, the quantal density of Is synapses was higher than that of Ib synapses (**Fig. 1C, D**). In both Ib and Is synapses, quantal density was significantly reduced in MN knockdowns of cac, Syt1, Rbp, unc-13 and VGluT (**Fig. 1C**). At Is synapses quantal density was also reduced in knockdowns of cac, Syt1, Rbp and unc-13, but not VGluT (**Fig. 1D**). We focused on three of the four MN knockdowns that caused the strongest reduction in quantal release at both Ib and Is synapses: cac, Rbp and unc-13.

**Figure 1.**
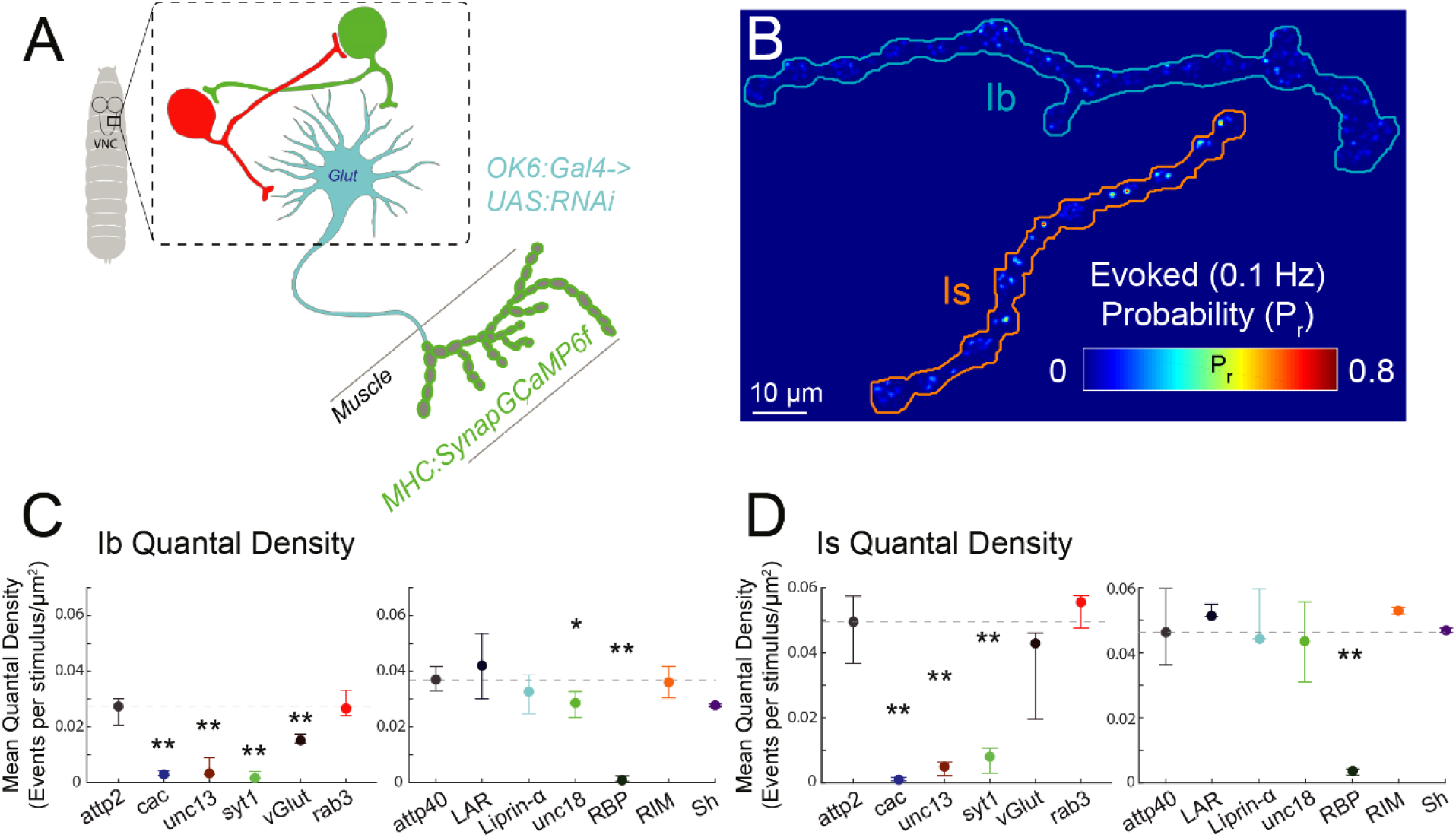
RNAi screen identifies presynaptic machinery knockdowns that weaken transmitter release by lb and Is MNs. (A) Overview of RNAi screen. OK6:Gal4 drives expression of RNAi in MNs. Quantal analysis of transmission via SynapGCaMP6f imaging of postsynaptic Ca2+ rise. (B) Representative *P,*heatmap calculated from failure analysis during train of 100 stimuli at 0.1 Hz. (C-D) Mean quanta! density (quantal content normalized to imaged NMJ area) for lb synapses (C) and Is synapses (D) from control animals (genotype) and from animals with **RNAi** of one of twelve genes encoding indicated presynaptic proteins *(Brp, Sh, cac, Syt1, rab3, RIM, Rbp, unc-13, unc-18, VG/uT, Liprin-a and LAR).* Points are median values for larvae and whiskers encompass the minimum and maximum values that are not outliers. Statistical comparisons Mann-Whitney test: *p < 0.05; **p < 0.01; ***p < 0.001.

### Effect on locomotion of perturbation of either the presynaptic release machinery or postsynaptic glutamate receptor

Considering that PHP only partially compensates for postsynaptic weakening and that *presynaptic* reduction in AP-evoked release triggers no compensatory change in *postsynaptic* quantal size (Newman, 2017; Genç and Davis, 2019), both classes of perturbation would be expected to result in a deficit in motor output. However, neither the postsynaptic perturbation, due to the GluRIIA^-/-^ mutant, nor the presynaptic RNAi knockdown of cac, Rbp or unc-13 in type I MNs with the motor-neuron specific *OK6-Gal4* line (Aberle et al., 2002; Sanyal, 2009), prevented larvae from surviving to adult, suggesting that neither disruption is too severe.

To examine the effect on motor behavior in detail, we tracked crawling activity of freely moving third-instar larvae (**Fig. 2A**). As described earlier (Günther et al., 2016; Lahiri et al., 2011; Loveless et al., 2019), larval locomotion alternated between two modes: active crawling and reorientation (**Movie S1**). The active crawling phase was characterized by peristaltic crawling that followed a relatively straight and persistent trajectory. Between periods of active crawling, there were phases of reorientation, where larvae ceased crawling, swept their heads from side to side, and then launched a new direction for forward movement.

**Figure 2.**
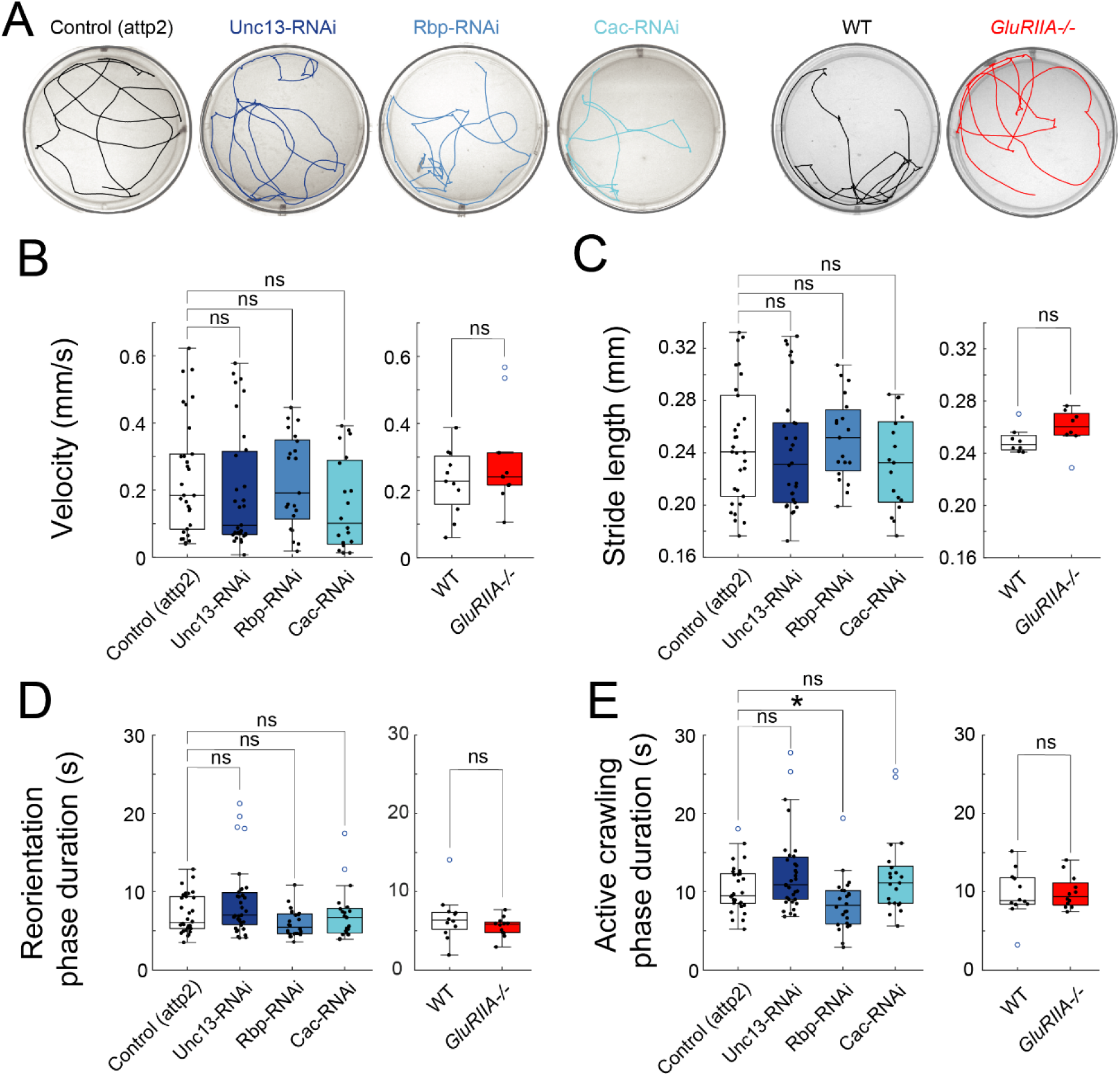
Crawling preserved despite impaired synaptic transmission. (A) Representative crawling tracks in 50mm wells. (B-E) Locomotion kinematics of crawling of animals with MN-specific **RNAi** of *unc13, Rbp* or cac, compared to Attp2 empty control (left), and GluRIIA-/- compared to WT control (right). (B) Mean velocity. (C) Mean stride length. (D) Mean duration in reorienting phase. (E) Mean duration of active crawling phase. Points are average value for each larva. Box plots depict median, the lower and upper quartiles, any outliers (open circles, computed using the interquartile range), and whiskers encompass the minimum and maximum values that are not outliers. Statistical comparisons Mann-Whitney test (B-E) *p<0.0.5; **p<0.01; ***p<0.001.

Strikingly, locomotion in both the presynaptic knockdowns and the GluRIIA*^-/-^* mutant was similar to that seen in control animals, with no change in average velocity or average stride length (**Figs. 2B, C and S2A**). We identified bouts of reorientation and measured their timing (**Fig. S2B**), but did not observe a change in the mean duration of reorientation bouts (**Fig. 2D**). The duration of the bouts of continuous crawling on a single trajectory were also indistinguishable between control and synaptic perturbations, except for a small (∼14%) reduction in one of the four perturbations, the presynaptic Rbp RNAi (**Fig. 2E**). In other words, the synaptic perturbations had little or no effect on core aspects of locomotion. Only when we quantified larval agility did we observe a statistically significant behavioral impact of the synaptic perturbations. By fitting a spline down the midline of the larva we were able to calculate body curvature (**Figure S2C, Supplemental Video 2**). We found a small but significant shift toward zero in the distribution of turning angles (angle between continuous trajectories) in the unc-13 and cac presynaptic RNAis and in the GluRIIA*^-/-^* animals, but not in the presynaptic Rbp RNAi (**Figure S2D**). The curvature distributions were significantly shifted to smaller curvatures in all three of the presynaptic perturbations and in the postsynaptic perturbation (**Figure S2E**), but these effects were small in magnitude.

Thus, despite the large reduction in AP-evoked transmission in the presynaptic knockdowns of cac, Rbp and unc-13, and the incomplete compensation for the reduction in postsynaptic strength caused by mutation of the GluRIIA subunit, larvae exhibited remarkably normal locomotor behavior while exhibiting only minor deficits in agility. We wondered if a change in MN firing pattern might compensate for reduced transmission by each individual AP. To examine this, we next studied natural neural activity in the intact, behaving animal.

### Increased duration of MN to muscle transmission bouts compensates for synaptic weakening

The *Drosophila* larva has 3 thoracic (T1-T3) and nine abdominal segments (A1-A9), most of which contain 30 bilateral body wall muscles. Each muscle is innervated by one of 26 different type Ib MNs, with a few muscles sharing a single Ib input, and the same 30 muscles innervated by two non-overlapping Is MNs, each of which innervates multiple muscles (Hoang and Chiba, 2001). The posterior-to-anterior (P-to-A) propagation of peristaltic muscle contraction simultaneously contracts the left and right sides of each segment to propel the larva forward (Gjorgjjeva 2013). To measure motor neuron input to the muscles while simultaneously observing forward-crawling peristaltic behavior, we turned to our restrained crawling setup, which consists of a linear chamber that restricts the larva to crawl in place while synaptic transmission is imaged in multiple segments (Newman 2017). Imaging of postsynaptic SynapGCaMP6f was used to measure the dynamics of synaptic transmission during crawling (**Figure 3A**).

**Figure 3.**
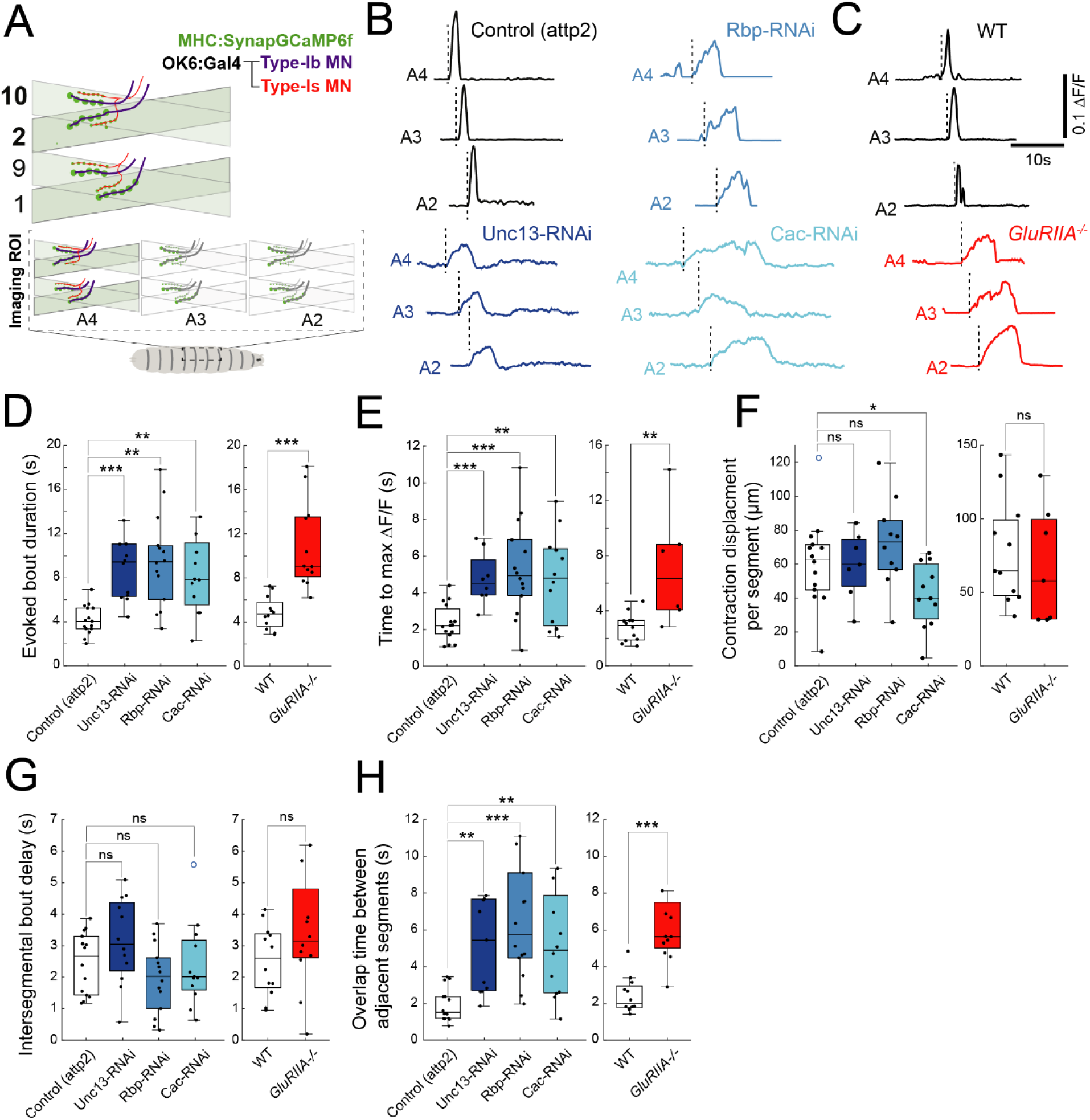
Pre- or post-synaptic weakening increases duration of synaptic transmission bouts during forward crawling waves. (A) Schematic of SynapGCaMP6f imaging of Ca^2^’ elevation due to influx through postsynaptic *GluR/1* receptor channels during MN-to-muscle synaptic transmission in intact, restrained larvae within segments A4, A3 and A2. (B, C) Representative l:i.F/F traces of SynapGCaMP6f in abdominal hemi-segments A2, A3 and A4 during P → A peristaltic wave for *unc-13, Rbp* or *cac* RNAis versus control (B) and G/uR//A-/-versus control (C). Start of bout marked by dashed vertical line. (D-H) lb MN-to-muscle 2 or 10 synaptic transmission dynamics in animals with MN-specific RNAi of *unc-13, Rbp* or *cac,* compared to attp2 empty control (left), and *G/uRIIA-1-*compared to WT control (right). (D) Average duration of activity bout in A2-A4. (E) Time to maximal l:i.F/F. (F) Contraction displacement per segment. (G) lnter-segmenlal delay from A4 lo A3 and A3 to A2. (H) Duration of overlap of bouts in A4 and A3, and in A3 and A2. Points are average value for each larva. Box plots depict median, the lower and upper quartiles, any outliers (open circles, computed using the interquartile range), and whiskers encompass the minimum and maximum values that are not outliers. Stalislical comparisons Mann-Whitney test (B-E) *p<0.0.5; **p<0.01; ***p<0.001.

Imaging from T1-A9 (**Movie S3**) revealed that, ∼95% of the time, when neighboring abdominal segments A2-A4 participate in a P-to-A wave, this is part of a peristaltic wave that travels along the entire length of the larva. To maximize the signal-to-noise of SynapGCaMP6f transmission imaging during peristaltic wave behavior, we focused on NMJs in dorsal A2-A4 segments (**Figures S3A**). As reported previously (Zarin 2019), we observed that the dorsal muscle cells of each segment contract together during P-to-A waves (**Movie S4**). Because muscle cells cluster into four co-activated muscle groups, we focused on Ib MNs innervating muscles 2 and 10 (**Figure 3A)**, which belong to the same peristaltic forward crawling group (Zarin 2019).

During propagation of P-to-A waves, each of the presynaptic release machinery KDs (RNAis of unc-13, Rbp and cac), as well as GluRIIA^-/-^, had an altered temporal pattern of synaptic transmission, with burst duration nearly doubling (**Figure 3B-D**) and the rise phase of the burst slowing (**Figures 3E and S3B**). Consistent with the preservation of total transmission, the amount of contraction displacement was similar to control animals in all but the cac RNAi (**Figure 3G**). The synaptic perturbations did not alter the speed of P-to-A wave propagation (**Figure 3H**). The RNAi and mutant larvae had greater temporal overlap between synaptic transmission bursts in adjacent segments (**Figure 3I**). These observations suggest that the nervous system preserves locomotion by increasing MN activity to compensate for the reduction in per AP MN to muscle synaptic transmission.

### Activity compensation involves a cell-autonomous increase in type Ib MN burst duration

Having observed overall activity compensation at presynaptically disrupted Ib NMJs, we wondered if such compensation also occurs at Is NMJs. To resolve the Is NMJs, which are smaller and whose SynapGCaMP6f signal is dimmer than the Ib NMJs, we focused on the NMJs of a single A1 segment including dorsal muscles 1, 2, 9 and 10 (**Figure 4A, Supplemental Video 5**). With our view limited to one segment, we were unable to distinguish activity bouts that were part of locomotory peristaltic waves from ones that are limited to A1 and thoracic segments, such as occur during head sweeping (Sawin-McCormack 1995; Humberg 2017). The integrated SynapGCaMP6f signal, which estimates the total synaptic transmission, were preserved at control levels in two of the three presynaptic perturbations (**Figure S4B**), while the degree of muscle contraction was preserved at control levels in all three presynaptic cases and the one postsynaptic cases of synapse perturbation (**Figure S4C**). A1 Ib NMJs approximately doubled bout duration and fraction time active in the unc-13, cac and RBP RNAi KDs, as well as in the GluRIIA null mutant (**Figure 4A-E**). Remarkably, in contrast to the changes seen in Ib synapses, no change was detected in the temporal pattern of synaptic transmission bouts in Is NMJs (**Figure 4F, G**).

**Figure 4.**
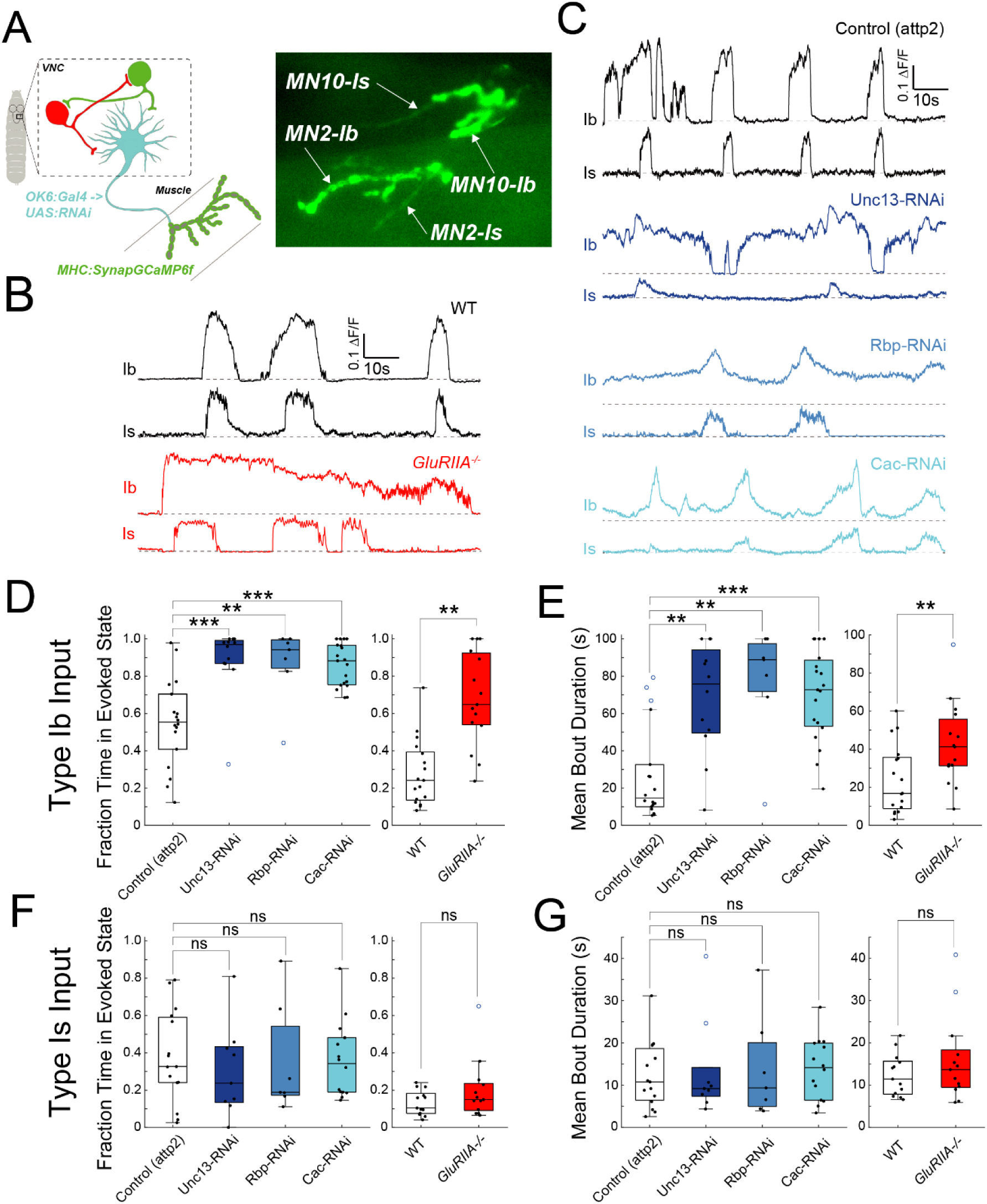
Increase MN burst activity is specific to lb inputs. (A) Schematic of SynapGCaMP6f imaging (left). Representative image of SynapGCaMP6 NMJ fluorescence at two lb inputs (MN2 and MN10). (B) Representative b.F/F traces for lb and Is in *G/uR//A-1·*and control. (C) Representative b.F/F traces for presynaptic RNAis and Attp2 control. (D) Fraction time in evoked state for lb. (E) Mean b.F/F bout durations for lb. (F) Fraction time in evoked state for Is. (G) Mean b.F/F bout durations for Is. Points are average value for each larva. Box plots depict median, the lower and upper quartiles, any outliers (open circles, computed using the interquartile range), and whiskers encompass the minimum and maximum values that are not outliers. Statistical comparisons Mann-Whitney test (B-E) *p<0.0.5; **p<0.01; ***p<0.001.

Cytosolic GCaMP6f recordings of *presynaptic* Ib MN activity from imaging of type Ib terminal axons revealed lengthened bursts of activity and fraction time active recorded during restrained locomotion in animals with weakened synaptic transmission (**Figure 5**). This indicates that increased bout duration and fraction time active of synaptic transmission results from increased MN firing.

**Figure 5.**
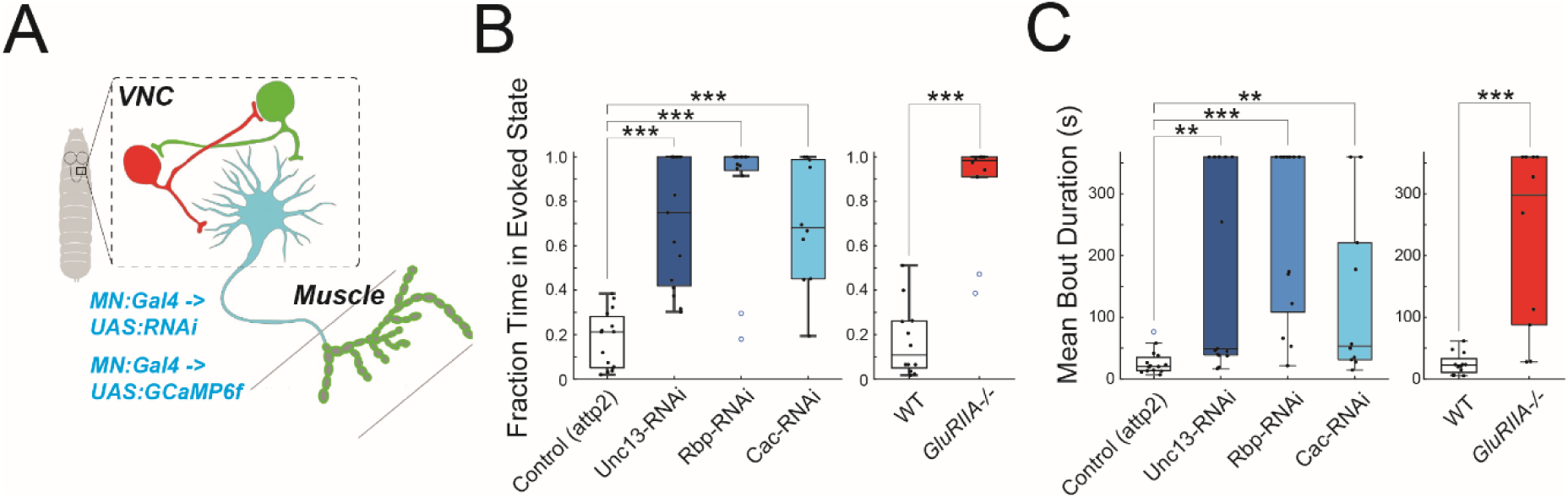
Effect of RNAis and *GluRIIN* on type lb MN presynaptic activity measured with MN GCaMP6f. (A) Schematic of imaging GCaMP6f in MNs. (B) Fraction time in evoked state. (C) Mean bout duration. Points are average value for each larva and bars show 95% confidence interval and median. Statistical comparisons Mann-Whitney test: *p < 0.05; **p < 0.01; ***p < 0.001.

### Altered activity in a central premotor input associated with MN firing homeostasis

The large increase that we observed in MN burst duration when synaptic transmission from MN to muscle was weakened led us to wonder if additional mechanisms, beyond increased excitability, contribute to increased type I MN activity. We considered that there may also be adjustments elsewhere in the locomotory circuit. We therefore turned our attention upstream of the MNs to the premotor neurons, particularly the inhibitory *period*-positive median segmental interneurons (PMSIs). The activity of PMSIs directly influences type I MN burst duration during locomotory waves, with elevated PMSI activity reducing type I MN burst duration (Kohsaka 2014), opposite to our observed compensation, and suggesting that reduced activity of PMSIs could also increase type I MN burst duration. We expressed cytosolic GCaMP6m in the PMSIs using R70C01-LexA, while we knocked down unc-13, Rbp or cac with UAS-RNAi exclusively in type I MNs with OK6-Gal4 (**Figure 6A).** We imaged activity in PMSI processes in the ventral nerve cord (VNC) across the abdominal segments, where they are clearly restricted to single segments (**Figure 6B**). In PMSIs of control (Attp2 empty) animals, we observed regular bouts of inhibitory activity that were uniform in amplitude and duration and traveled in both the P→A and A→P directions (**Figure 6C, Supplemental Video 5**). Inhibitory activity bout duration in the PMSIs did not change, nor did bout amplitude, with the exception of a small effect in the Rbp knockdown (**Figure S5A, B**). In addition, there was no effect on the intersegmental delay between the onset of bouts during peristaltic waves (**Figure S5C**). However, in the knockdowns of unc-13, Rbp or cac there were dramatic effects on both the frequency of peristaltic waves and activity bouts, as well as the fraction active time—all of which decreased substantially (**Figure 6C-F**). These observations suggest that type I MN increased firing in animals with weakened MN to muscle synaptic transmission involves an upstream circuit adjustment that decreases inhibitory drive to type I MNs.

**Figure 6.**
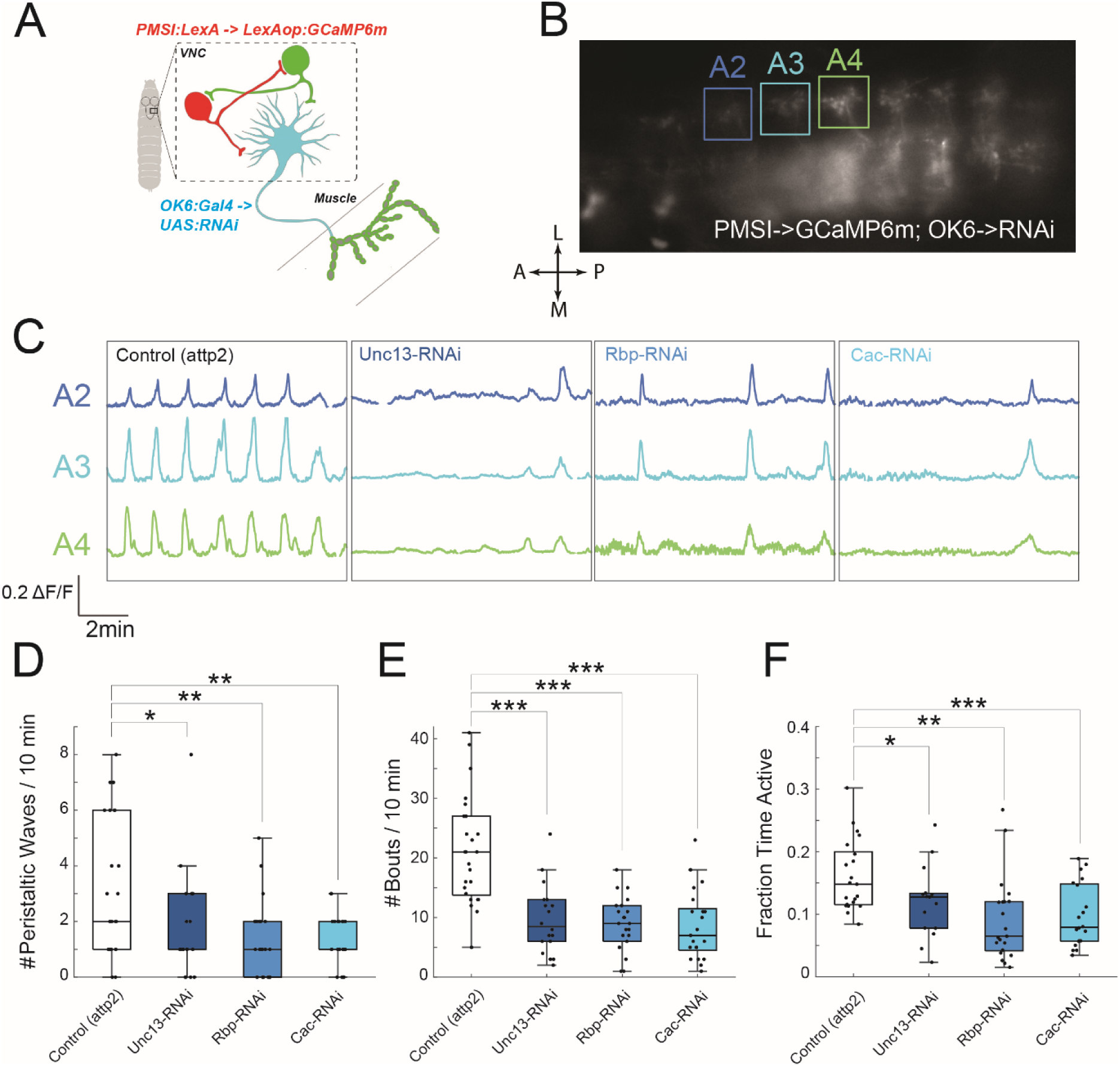
Weakening of MN to muscle synapse decreases activity of PMSI inhibitory pre-motor neurons. (A, B) Schematic (A) and image (B) of cytosolic GCaMP6m *(R70C01-LexA)* activity representing a P→A peristaltic wave in PMSls. (C) Representative GCaMP6f l’IF/F traces in PMSI of segmentsA2, A3 and A4 traces in animals with MN-specific RNAi for *unc13, rbp* or *cac,* compared to Attp2 empty control. (D) Number of peristaltic waves per 10 minutes. (E) Mean number of b.F/F activity bouts per 10 minutes. (F) Fraction time active. Box plots depict median, the lower and upper quartiles, any outliers (open circles, computed using the interquartile range), and whiskers encompass the minimum and maximum values that are not outliers. Statistical comparisons Mann-Whitney test (B-E) *p<0.0.5; **p<0.01; ***p<0.001.

### Timing of homeostatic firing adjustment in Ib MNs

PHP at the *Drosophila* larval NMJ begins to boost AP-evoked transmitter release within minutes after onset of block of postsynaptic iGluRs by the wasp venom *Philanthotoxin*-433 (PhTox) (Frank 2006). We wondered whether the increase in Ib MN activity, which we observe in synaptic weakening, also occurs quickly. Since PhTox cannot be readily introduced into the intact larva and reduced expression of GluRIIA occurs slowly following a genetic manipulation, we turned to the temperature-sensitive *Drosophila* dynamin mutant, *shibire*^ts1^ (*shi*^ts1^), which reduces transmitter release at the restrictive temperature within minutes, due to block of vesicle recycling (Koenig 1989). We expressed UAS-*shi*^ts1^ in type I MNs under the OK6 MN-Gal4. Initially, we held larvae at a permissive temperature (18°C) and then switched to a restrictive temperature (26°C) and imaged the SynapCGaMP6f reporter of synaptic transmission in segments A2-A4 during P→A peristaltic waves over 30 min, which we divided into three successive 10 min bins following transition to the restrictive temperature. We compared *shi*^ts1^-expressing animals (UAS-*shi*^ts1^; OK6-Gal4) to controls (UAS-*shi*^ts1^) that lacked the Gal4 driver (**Figure 7**). At the restrictive temperature, synaptic transmission ΔF/F bout amplitude was reduced in the *shi*^ts1^ expressing larvae compared to control by the 10-20 min bin (**Figure 7A**). Synaptic transmission bout duration increased 10 minutes later, in the 20-30 min bin (**Figure 7B**). Thus, the homeostatic change in MN activity pattern initiates over a period of ∼10 minutes following acute presynaptic reduction in glutamate release.

**Figure 7.**
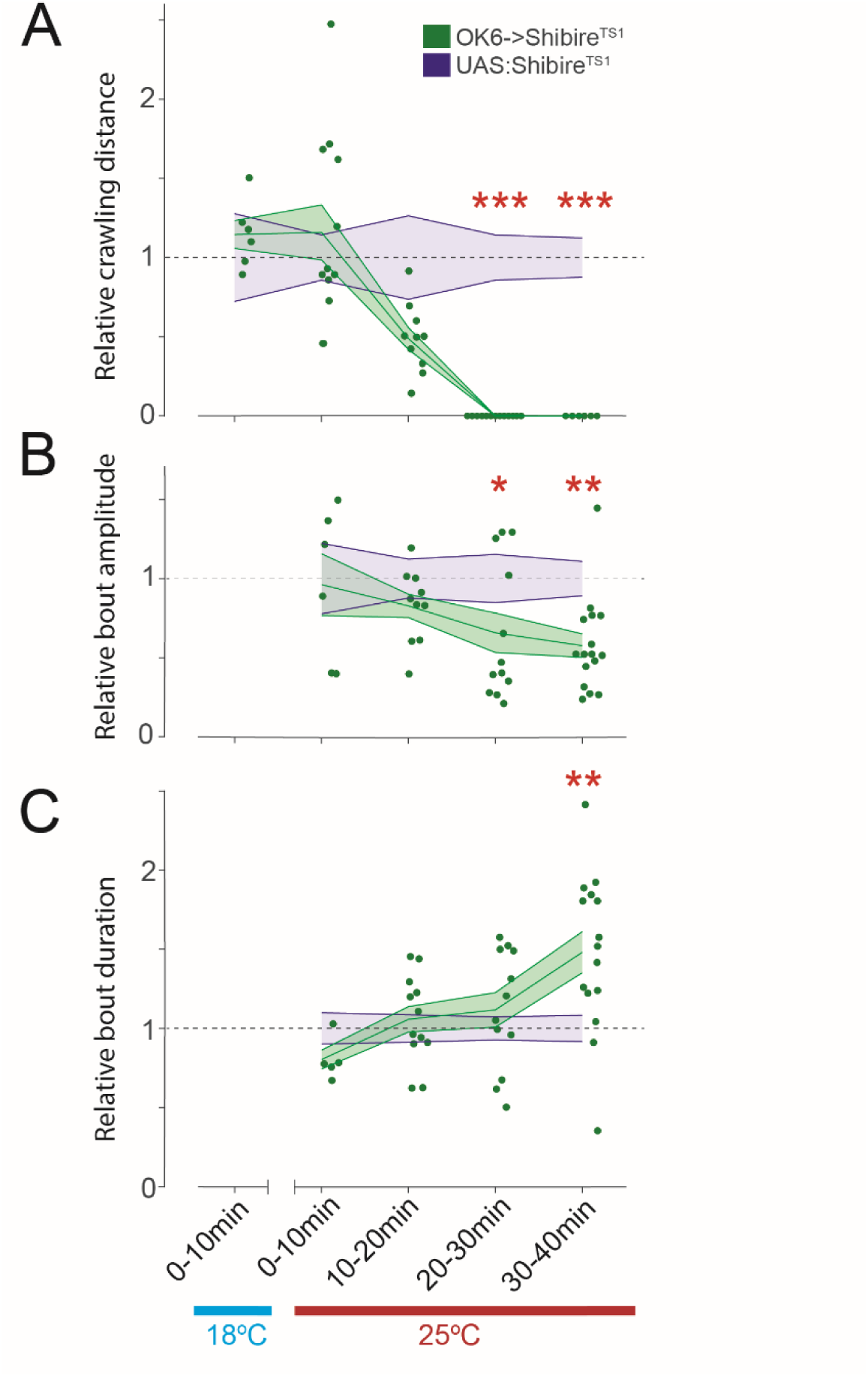
Time course of bout duration increase versus Shibirers’-induced decrease in synaptic transmission. (A) Relative crawling distance due to OK6->Shibirers, activation over time. (B) Relative ll.F/F bout amplitude over time. (C) Relative t:.F/F bout duration over time. Points are average value for each larva. Error clouds represent S.E.M. Statistical comparisons Mann-Whitney test: *p < 0.05; **p < 0.01; ***p < 0.001.

## Discussion

Homeostatic mechanisms in the nervous system maintain a balance between excitation and inhibition and ensure circuit throughput by adjusting synaptic transmission and excitability following activity-dependent plasticity changes and pathological perturbations (Marder 2006; Turrigiano 2012; Davis and Muller, 2015 and others by Turrigiano; Kulik et al., 2019). Presynaptic homeostatic plasticity (PHP), a key homeostatic mechanism studied extensively in the Drosophila larval NMJ, has been shown to compensate for postsynaptic reduction in sensitivity to transmitter by increasing transmitter release *via* a retrograde signaling system (Peterson, 1997; Davis, 2015; Frank 2006). However, PHP only occurs in one of the two excitatory glutamatergic inputs to muscle, the type Ib MN, and even there only partially compensates with increased AP-evoked release probability (P_r_) for the reduction in the amplitude of the postsynaptic quantal response (Newman, 2017; Genç and Davis, 2019). Despite this synaptic shortfall, we find that postsynaptically weakened larvae have close to normal locomotion under non-challenging conditions, right side up on a substrate. Moreover, severe presynaptic reduction in glutamate release, elicited by knockdown of key components of the MN presynaptic transmitter release machinery, also leaves locomotion near normal under this natural condition. This suggests that additional forms of homeostatic adjustment may be engaged in parallel to fully compensate for reduced synaptic transmission per AP.

We find that larvae with weakened neuro-muscular transmission *increase* the duration of locomotory MN bursts and the total fraction of time active during both locomotory waves and non-locomotory movement. This adjustment occurs in tonic type Ib MN 1, 2, 9 and 10, but not in the phasic Is MN (MSNISN) that innervates the same muscles (Kohsaka 2014; Zarin 2019). Crawling velocity has been shown to be regulated by peristaltic wave frequency rather than stride length (Heckscher 2012). In accord with this, we observe that peristalsis wave speed and frequency are preserved in both the pre- and postsynaptically weakened larvae, consistent with the observed maintenance of normal crawling velocity. The activity adjustment increases the amount of time that neighboring segments are simultaneous active, which may help to generate the same contractile displacement that individual segments normally exert. The increase Ib transmission bout duration offsets the reduction in magnitude of postsynaptic Ca^2+^ influx through the GluRII receptor channels, suggesting that the homeostatic mechanism may operate to maintain a setpoint of integrated intracellular Ca^2+^ at the postsynaptic density.

The increase in MN firing duration may be due to a change in the activity of PMSIs, which provide an inhibitory central premotor neuron input to type I MNs. We observe that, in synaptically weakened animals, locomotion-associated burst activity is *decreased* in PMSIs. These results agree with earlier work that showed that the opposite effect optogenetic stimulation that *increases* the activity of PMSIs has the opposite effect: shortening MN burst duration (Kohsaka 2014). Interestingly, PMSIs innervate type Ib MN1, 2, 9 and 10, but not the Is MSNISN that converges on the same muscles (Zarin 2019). The selectivity of innervation of type Ib MNs is consistent with the selective firing adjustment in Ib MNs and lack of such adjustment in the type Is MN. Thus, weakening of synaptic transmission from type Ib and Is MNs to muscle--*i.e.* the output end of the locomotory system—triggers a two-pronged change that increases type Ib MN firing: a cell-autonomous increase in type Ib MN excitability and a circuit adjustment of inhibitory inputs to type Ib MNs.

Type Ib inputs have low P_r_ synapses that tend to facilitate, and their activity ramps up and down slowly, whereas type Is inputs have high P_r_ synapses that tend to depress, and their activity turns on and off abruptly so that they burst only during the maximal contraction phase (Newman, 2017; Akbergenova 2018; Aponte-Santiago 2020). Seeing that both PHP and firing homeostasis occur in Ib but not Is MNs, the Is input provides constancy that can ensure proper locomotor wave dynamics, while the Ib input provides plasticity to adjust to changing conditions, between them ensuring robust locomotor behavior. Reduced inhibitory drive from pre-MNs onto the Ibs is expected to further contribute to increased Ib activity (**Fig. 8**). Firing homeostasis and PHP are complementary and multiplicative, with firing homeostasis increasing the number of APs and PHP increasing the amount of transmitter release per AP.

**Figure 8.**
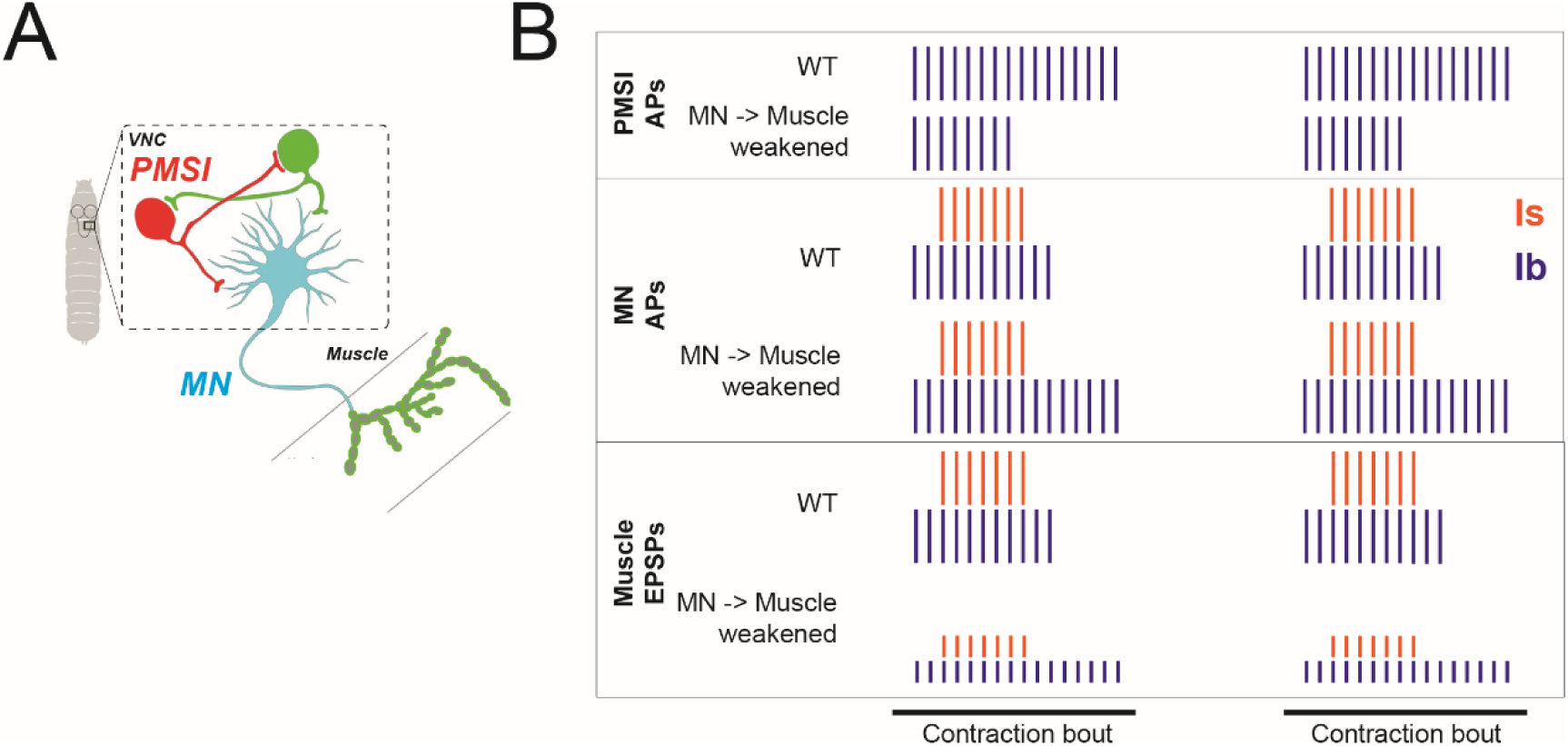
Model depicting two-forms of homeostatic compensation in response to weakening of synaptic transmission from MN to muscle. (A) Schematic of inhibitory pre-motor neurons synapsing onto Type I motor neurons (MN) (inhibitory PMSI, red; excitatory cell, green). (B) In response to weakened transmission from the MNs onto the muscle (lower) there is a compensatory increase in duration of Type lb firing, but not Is (middle), in addition to a decreased burst duration of inhibitory PMSI premolar neurons (upper).

## Experimental Model and Subject Details

### Drosophila

Transgenic flies were made using standard germline transformation by embryo injection (BestGene, Chino Hills CA). 24B-Gal4 (Luo et al., 1994), GluRIIASP16 and Df(2L)clh4 (Petersen et al., 1997) flies were from Corey Goodman (UC Berkeley). UAS-Shibire^TS1^ (Kitamoto, 2001) was from Kristen Scott (UC Berkeley). OK6-Gal4 (#64199), R70C01-LexA (#54927), UAS-GCaMP6f, UAS-GCaMP6m, LexAop-GCaMP6m, UAS-Dicer2, and UAS-RNAi (TRiP) lines using either the attp2 or attp40 docking sites were from Bloomington Drosophila Stock Center (Bloomington, IN). Flies were raised on standard corn meal and molasses media at 25°C, except those with RNAis, which were raised at 28°C, or those used in Shibire^TS1^ experiments, which were raised at 18°C. Both male and female wandering third instar larvae were used in all experiments. When required, third instar larvae were screened using an Axio Zoom.V16 microscope (Carl Zeiss, Oberkochen, Germany) through the use of balancers with larval markers including CyOGFP (3xP3-EGFP variant) and TM6B. Only actively crawling larvae were used for experiments.

The following genotypes were used:

Control (attp2) (UAS-Dicer2;OK6-Gal4/+; SynapGCaMP6f/attp2 (empty)),

cac (UAS-Dicer2;OK6-Gal4/+;SynapGCaMP6f/UAS-cac-RNAi (#27244),

unc13 (UAS-Dicer2;OK6-Gal4/+;SynapGCaMP6f/UAS-unc13-RNAi (#29548),

syt1 (UAS-Dicer2;OK6-Gal4/+;SynapGCaMP6f/UAS-syt1-RNAi (#28508),

unc13 (UAS-Dicer2;OK6-Gal4/+;SynapGCaMP6f/UAS-unc13-RNAi (#27538),

vGlut (UAS-Dicer2;OK6-Gal4/+;SynapGCaMP6f/UAS-vGlut-RNAi (#27538),

rab3 (UAS-Dicer2;OK6-Gal4/+;SynapGCaMP6f/UAS-rab3-RNAi (#25953),

Control (attp40) (UAS-Dicer2;OK6-Gal4/attp40 (empty);SynapGCaMP6f/+),

LAR (UAS-Dicer2;OK6-Gal4/UAS-LAR-RNAi (#40938);SynapGCaMP6f/+),

Liprin-α (UAS-Dicer2;OK6-Gal4/UAS-Liprin-α-RNAi (#28301);SynapGCaMP6f/+),

unc18 (UAS-Dicer2;OK6-Gal4/UAS-unc18-RNAi (#51925);SynapGCaMP6f/+),

RBP in attp40 (UAS-Dicer2;OK6-Gal4/UAS-RBP-RNAi (#54828);SynapGCaMP6f/+),

RBP in attp2 (UAS-Dicer2;OK6-Gal4/+;SynapGCaMP6f/UAS-RBP-RNAi (#29312)),

RIM (UAS-Dicer2;OK6-Gal4/UAS-RIM-RNAi (#55741);SynapGCaMP6f/+),

Sh (UAS-Dicer2;OK6-Gal4/UAS-Sh-RNAi (#55347);SynapGCaMP6f/+),

wild-type control (w1118;+;+), wild-type SynapGCaMP6f (WT; w1118;+/+;SynapGCaMP6f/+),

GluRIIA (GluRIIASP16/Df(2L)clh4;SynapGCaMP6f/+),

OK6->Shibire^TS1^ (w1118;OK6-Gal4/+;SynapGCaMP6f/UAS-Shibire^TS1^),

UAS: Shibire^TS1^ (w1118;+;SynapGCaMP6f/UAS-Shibire^TS1^,

PMSI Control (UAS-Dicer2; R70C01-LexA/OK6-Gal4; attp2 (empty)/LexAop-GCaMP6m),

PMSI Unc13-RNAi (UAS-Dicer2; R70C01-LexA/OK6-Gal4;UAS-Unc13-RNAi/LexAop-GCaMP6m),

PMSI RBP-RNAi (UAS-Dicer2; R70C01-LexA/OK6-Gal4;UAS-RBP-RNAi/LexAop-GCaMP6m),

PMSI cac-RNAi (UAS-Dicer2; R70C01-LexA/OK6-Gal4;UAS-cac-RNAi/LexAop-GCaMP6m).

### Method Details

#### Larval Locomotion Assay

Multiplexed tracking of crawling larvae was accomplished by outfitting a 6-well plate filled with 2% agarose gel leaving. A gap of approximately 3 mm was left between the surface of the agarose and the cusp of the well to minimize the movement of larvae in the z-direction. A single larva was placed in each well of the plate for tracking individual animals. Larvae were filmed crawling for a single 10 minute trial at a rate of 10 fps backlit with an IR LED array. The imaging chamber was kept at 21 °C and illuminated with a 60W incandescent bulb.

#### SynapGCaMP6f Optical Quantal Imaging

For high-resolution quantal analysis, third instar larvae were initially dissected in HL3 solution containing (in mM): 70 NaCl, 5 KCl, 0.45 CaCl2·2H2O, 20 MgCl2·6H2O, 10 NaHCO3, 5 trehalose, 115 sucrose, and 5 HEPES, with pH adjusted to 7.2. After removing the brain, larval fillets were washed and imaged in HL3 solution containing 1.5 mM Ca2+ and 25 mM Mg2+, unless specified otherwise (refer to Figures S3G and S5S–S5V). Fluorescence images were captured using a Vivo Spinning Disk Confocal microscope (3i Intelligent Imaging Innovations, Denver, CO), equipped with a 63x 1.0NA water immersion objective (Zeiss), a 488nm (50 mW) LaserStack laser, a Yokogawa CSU-X1 A1 spinning disk (Tokyo, Japan), a standard GFP filter, and an EMCCD camera (Photometrics Evolve, Tucson, AZ). Unless noted otherwise (see Figure S7F and S7G, where a 20x water objective was used), all fillet recordings were performed on ventral longitudinal abdominal muscle 4 at segments A2-A5 of third instar larvae. Fields of view were chosen to maximize the area of both Ib and Is terminals within a single focal plane. The imaging area was then cropped to the NMJ area.

Nerve stimulation was conducted using a suction electrode connected to a Stimulus Isolation Unit (SIU, ISO-Flex, A.M.P.I Jerusalem, Israel) with a 75-100 μs stimulus duration. The stimulation intensity was adjusted to recruit both Ib and Is axons, verified during imaging. Nerve stimulation and imaging were synchronized using custom MATLAB (MATLAB Version R2015b and Image Processing Toolbox Release R2015b, MathWorks, Natick, MA) scripts to control the SIU and trigger imaging episodes within SlideBook (v6.0.9, 3i Intelligent Imaging Innovations). Episodic stimulation and imaging involved acquiring a series of 10 images (50 ms exposures) per episode. Each episode included at least 3-4 baseline frames before nerve stimulation. 200 stimulus trials at 0.1 Hz were used for analysis to minimize statistical fluctuations’ effects on the calculation of Pr at individual synapses (see below). Stimuli were time-locked to the start of the recording to ensure stimuli registration.

#### In vivo Intact Larval SynapGCaMP6f Imaging

In vivo imaging of intact larvae was performed similarly as in Newman et. al. 2017. For in vivo imaging, third instar larvae were placed in custom-made gas-permeable poly(dimethylsiloxane) (PDMS). For imaging, larvae were mounted in the PDMS chamber, which had an internal depth of 500 μm and width of 1 mm. The larvae were oriented with their dorsal surface facing up and sealed with a coverslip. Imaging was performed using an Axio Zoom.V16 microscope with a 2.3x0.57 NA objective, a FS 38 HE filter set (with excitation at 470/40 nm and emission at 525/50 nm), and a Teledyne Prime 95B camera. Images were acquired continuously for either 6 or 10 minutes at a frequency of 20 Hz, with final magnifications of either 80x or 260x and a 2 μm depth of field to keep NMJs in focus during slight movements. High-power (260x) imaging data were collected from dorsal muscles 1, 2, 9 and 10 in either segment A1 or A2, whereas low-power (80x) imaging captured SynapGCaMP6f activity from segments A2, A3, and A4, where background autofluorescence from the gut and other organs minimally interfered with SynapGCaMP6f signal. The anterior segments provided clearer images with less autofluorescence and light scattering. VNC imaging was performed at the low-power magnification and encompassed the entire VNC. Only larvae showing sustained activity were included in the analysis, while those with excessive movement or focus issues were excluded.

### Quantification and Statistical Analysis

#### Optical Quantal Image Analysis

Quantal image analysis was performed using techniques described in previous work (Peled, 2014; Newman 2017). In summary, optical quantal image analysis was conducted using custom MATLAB routines. The process involved filtering images to reduce noise, separating them into Ib and Is regions based on fluorescence, and correcting for motion and bleaching. Stimulus trials were excluded if they were out of focus, moved, or the stimulus failed.

For automatic detection of ΔF spots, templates were created from manually identified spots. ΔF response images were generated, and participating pixels were identified based on correlation with the template response. Release probability maps were constructed by identifying ΔF spot centers across multiple trials and color-coded to show the frequency of spot centers.

Individual release sites were manually identified, and Pr values were calculated by dividing the number of ΔF spot centers by the number of trials. Quantal density, an unbiased metric, was calculated by averaging the number of ΔF spot centers over all trials and normalizing by the imaged area.

#### Behavioral Analysis

Movies were processed with custom Matlab routines to assess two main features of larval locomotion, bouts of crawling along a continuous trajectory and reorientation events, as described previously (Günther et al., 2016). Trajectories of each larvae were calculated with a 2D centroid tracking algorithm. Centroids within 1 mm of each other across a 2 second window were merged by calculating the center of mass value, reducing noise in the crawling trajectory.

Reorientation events were scored by fitting a 3rd degree polynomial along the length of the larva and calculating the curvature. Curvature values were low-pass filtered, and a reorientation event was scored when curvature exceeded the middle quartile value calculated across the entire movie for a duration of at least 2 s. Total distance traveled was determined from the length of the trajectory over the course of the experiment. Time spent during reorientation events was calculated from the number of centroids used to calculate a smoothed point scored as a reorientation event. Time spent crawling on a continuous trajectory was calculated from the time in between reorientation events, excluding periods where the larva stopped crawling.

#### In vivo Image Analysis

Image analysis was performed similar to Newman et. al 2017. In brief, all image analysis was conducted using custom MATLAB routines. To stabilize the moving and contracting NMJs in vivo, the movies underwent a series of image registration steps. Each recording typically included multiple Ib and Is NMJ pairs, which were processed individually. For high-power imaging (260x magnification) of the NMJ using either muscle-driven SynapGCaMP6f or motor-neuron-driven GCaMP6f, initial corrections for coarse, rigid translational movements were achieved by calculating x- and y-displacements to maximize the 2D cross-correlation between successive frames. Subsequent registration of the NMJs within the tracked areas involved affine and diffeomorphic nonlinear (Demon) transformations (MATLAB Image Processing Toolbox Release R2015b). A high-intensity frame, featuring both active Ib and Is NMJs, served as the reference for these corrections, which adjusted for local shape changes during muscle contractions. Fluorescence data were obtained from manually defined regions of interest (ROIs) around the NMJs or cell-bodies in the VNC. For lower-power (80x) and VNC imaging, only the translation and affine registration steps were performed.

To isolate the synaptic fluorescence changes, spatially uniform muscle fluorescence signals (Fm) and any background fluorescence changes were subtracted from the absolute fluorescence prior to calculating Ib and Is ΔF/F. This was done by calculating the mean fluorescence intensity from a region adjacent to the NMJs but free of synapses. Although bleaching was minimal under these imaging conditions, subtracting Fm also helped correct for any slow bleaching that occurred during prolonged recordings. This method ensured that the synaptic component of the SynapGCaMP6f or GCaMP6f fluorescence changes was accurately isolated in the complex in vivo environment of a behaving larva.

For either NMJ or VNC imaging, the onsets of evoked activity bouts were defined when either SynapCaMP6f or GCaMP ΔF/F surpassed 10% of the baseline level, whereas the ends of bouts fell below this threshold. Evoked activity bouts were manually verified by examining individual movies. Bout durations were calculated using the length in seconds of each activity bout, and the time to max was calculated from the start of the bout to maximum ΔF/F of the bout. The contraction displacement per segment was measured from the degree of translation from the center-of-mass of the NMJ being registered. Intersegmental bout delay is calculated from time of onset of one bout in a segment to the onset of the successive bout in the neighboring segment during P → A peristaltic waves. Overlap time between adjacent segments was measured when both NMJs from two neighboring segments were simultaneously active. Mean bout amplitude was calculated from measuring the average ΔF/F of the entire bouts. For high-power imaging (260x), the calculations for integrated fluorescence and muscle contraction magnitude are described in detail in Newman et. al. 2017. Fraction-time in evoked state is calculated by dividing the total time of evoked activity by the total time of the acquired movie.

## Code availability

Code will be available in a Github repository upon publication.

## Data availability

Source data will be provided with this paper upon publication.

## Acknowledgements

We thank William Liu and Susan Younger for help with fly genetics and all of the members of the Isacoff lab for helpful discussion. The study was supported by the National Institutes of Health (R01NS107506 to E.Y.I.). E.Y.I. is a Weill Neurohub Investigator.

## Contributions

A.H. designed and performed the experiments and analysis on locomotion, imaging in restrained larvae of activity of type Ib MNs and PMSIs and synaptic transmission imaging by type Ib MNs. Z.L.N. designed and performed the optical quantal analysis RNAi screen that identified Unc-13, Rbp and Cac knockdowns as severe perturbations of glutamate release. A.H. and E.Y.I. conceived and designed the project and co-wrote the paper with input from all of the authors. E.Y.I. oversaw the study.

