## Supplemental Figures for "Circuit firing homeostasis following synaptic perturbation ensures robust behavior"

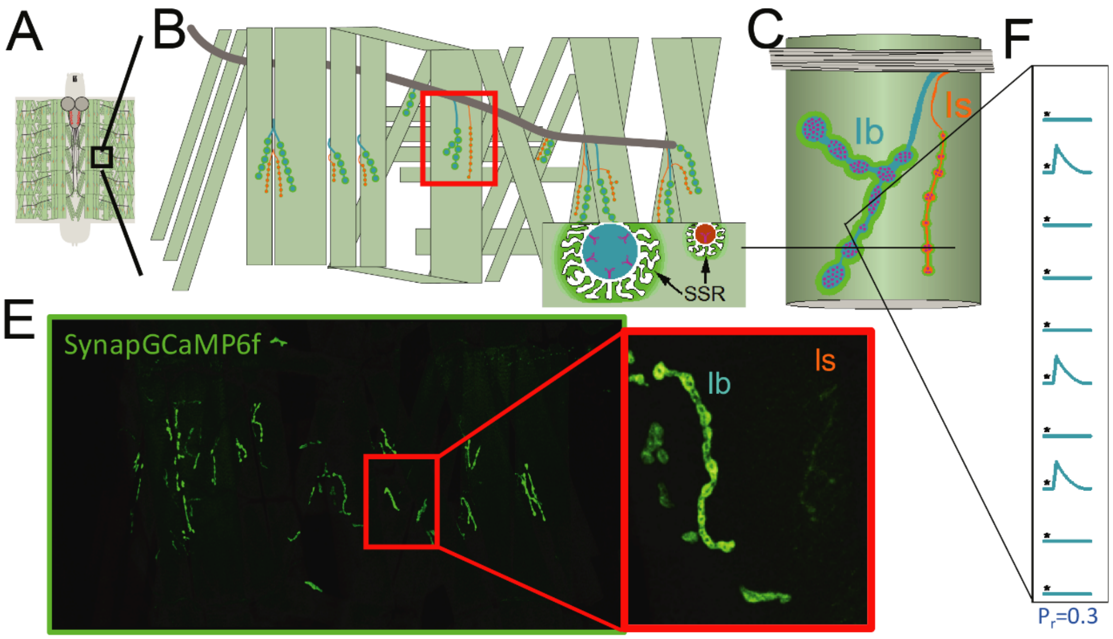


**Figure S1, related to Figure 1. Optical quantal analysis of glutamatergic transmission from type I motor neurons at the Drosophila larval NMJ.** (A-D) Cartoons of Drosophila larval filet preparation. (B, C, D) Increasing magnitude blowups of single abdominal segment (black dashed ROI in (A)), showing type Ib (blue) and type Is (orange) motor neurons innervating body wall muscles, surrounded by postsynaptic SynapGCaMP6f (green) in the muscle, which concentrates in the subsynaptic reticulum (SSR in (B)) through interaction with the PDZ protein Discslarge (Dlg). (E) Fluorescent image showing region depicted in (B) with blowup on right of ROI shown in red square. Type Ib postsynaptic areas are highly enriched in Dlg and therefore in SynapG-CaMP6f, while Type Is postsynaptic areas have less Dlg and so are dim. (F) Cartoon depiction of fluorescence responses in one postsynapse to 10 presynaptic stimuli given at 0.1 Hz, shows 3 responses, indicating a probability of action potential evoked release (Pr) of 0.3.

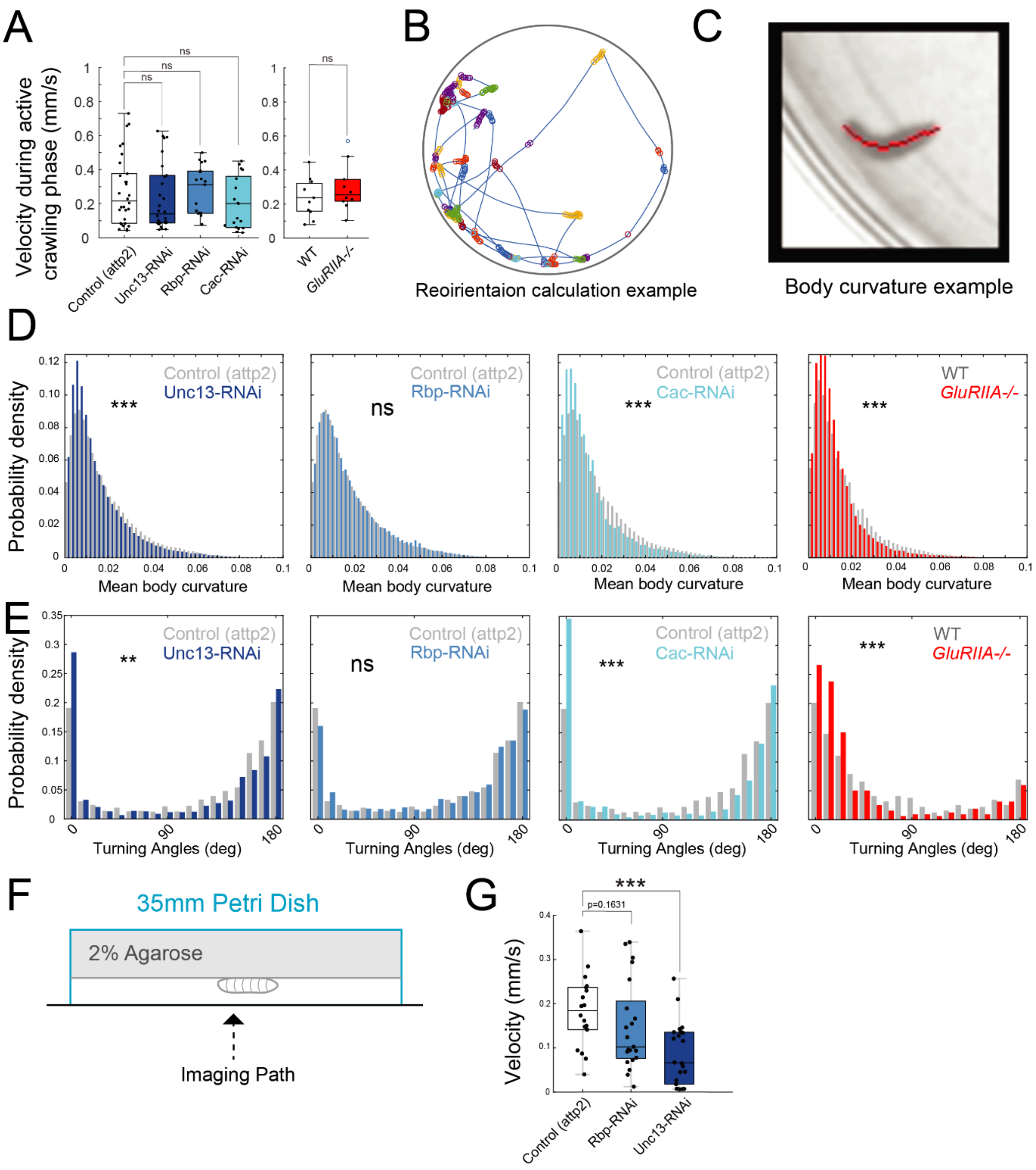


**Figure S2, related to Figure 2. Effect on crawling agility of perturbation of presynaptic release machinery or postsynaptic receptor.** (A) Velocity during the active crawling phases (B) Example of active crawling phases (blue lines) punctuated by reorientation phases (circles) (C) Representative spline (red) fit along the body axis of the larva (D) Probability densities of mean body curvatures for RNAis and GluRIIA mutant (E) Distributions of turning angles from 0 to 180 degrees (F) Diagram of challenged, upside-down crawling assay on 2% agarose plate. (G) Velocity of larvae in the upside-down crawling assay. Points are average value for each larva. Box plots depict median, the lower and upper quartiles, any outliers (open circles, computed using the interquartile range), and whiskers encompass the minimum and maximum values that are not outliers. Statistical comparisons Mann-Whitney test (B-E) *p<0.0.5; **p<0.01; ***p<0.001.

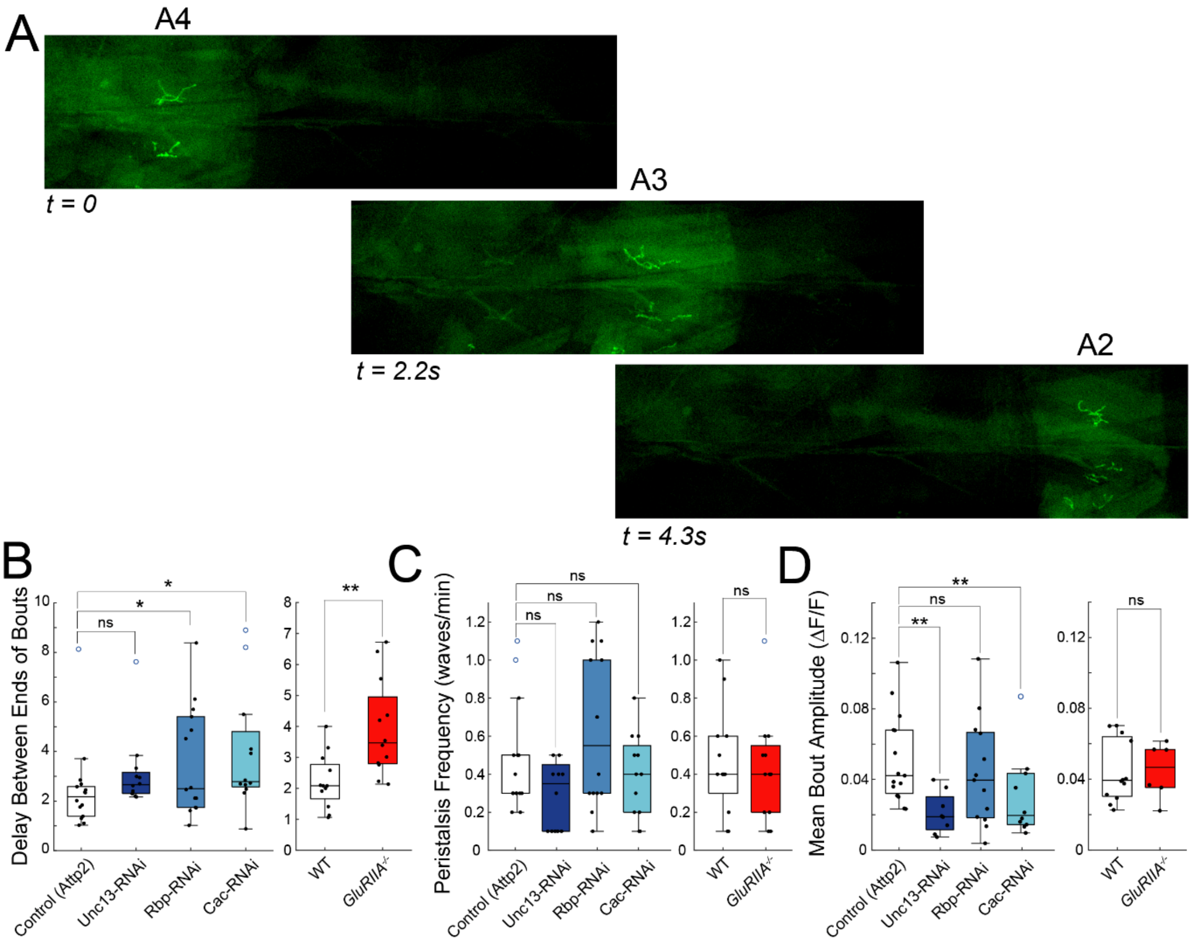


**Figure S3, related to Figure 3. Extended properties of posterior-to-anterior (P → A) peristaltic bouts.** (A) Representative P → A wave from control movie (20 fps). (B) Average delay between ends of bouts in neighboring segments during P→A peristaltic waves. (C) P → A peristaltic wave frequency. (D) P → A mean bout amplitude. (B-D) Points are average value for each larva. Box plots depict median, the lower and upper quartiles, any outliers (open circles, computed using the interquartile range), and whiskers encompass the minimum and maximum values that are not outliers. Statistical comparisons Mann-Whitney test (B-D) *p<0.0.5; **p<0.01; ***p<0.001.

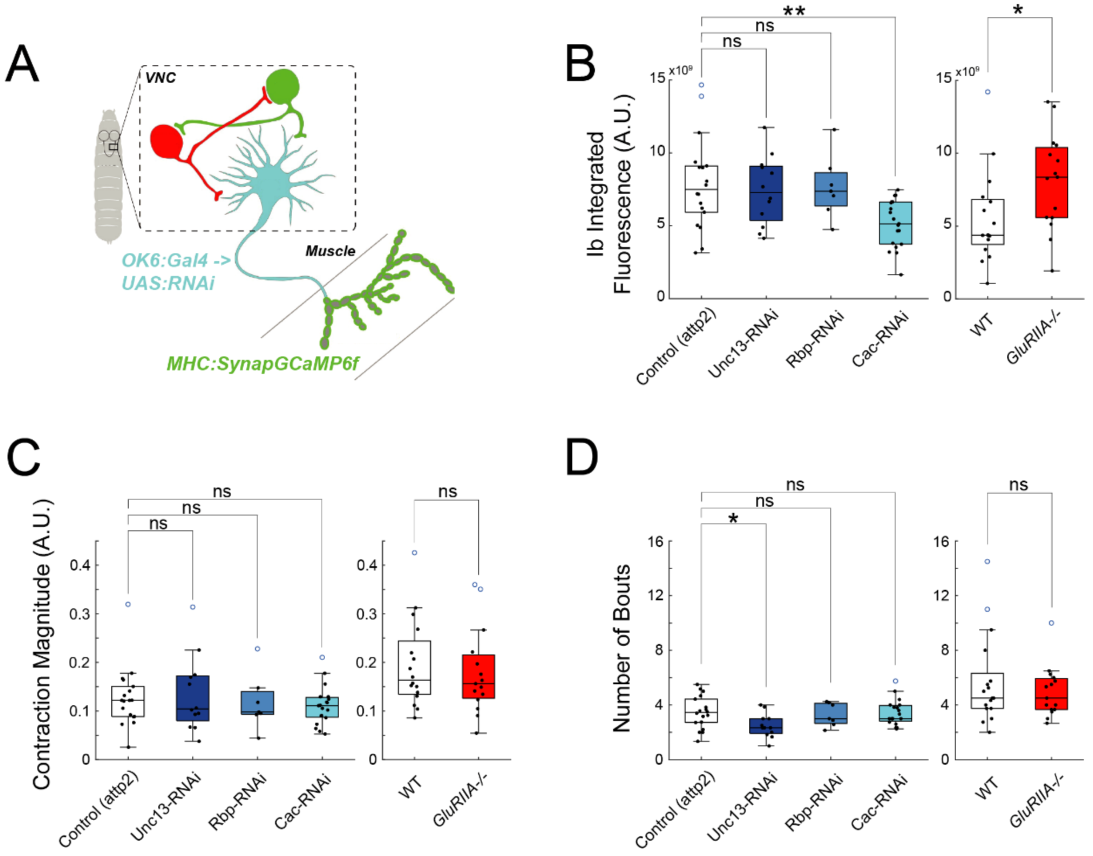


**Figure S4, related to Figure 4. Extended properties of Ib and Is inputs in segment A1.** (A) Schematic of SynapGCaMP6f imaging. (B) Total MN synaptic transmission measured with postsynaptic SynapGCaMP6f integrated fluorescence. (C) Contraction magnitude as represented by the determinant of the affine transmformation used in image registration. (D) Number of bouts over duration of recording (6 minutes). Points are average value for each larva. Box plots depict median, the lower and upper quartiles, any outliers (open circles, computed using the interquartile range), and whiskers encompass the minimum and maximum values that are not outliers. Statistical comparisons Mann-Whitney test (B-E) *p<0.0.5; **p<0.01; ***p<0.001.

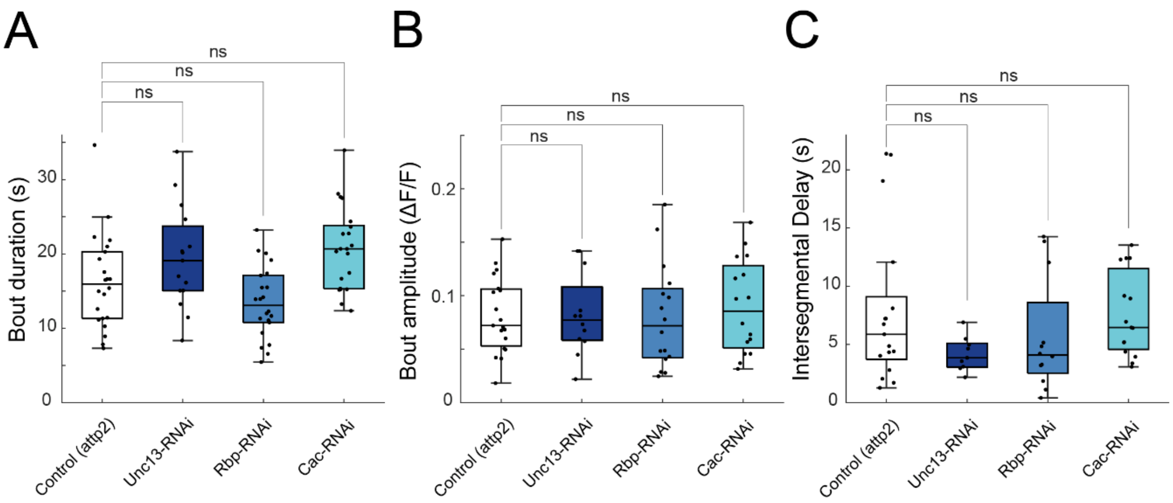


**Figure S5, related to Figure 6. Extended results for effect of weakened MN to muscle synapse on activity of PMSI inhibitory pre-motor neurons.** (A) Bout duration. (B) Bout amplitude. (C) Intersegmental delay between bout onset during P → A peristaltic waves. Box plots depict median, the lower and upper quartiles, any outliers (open circles, computed using the interquartile range), and whiskers encompass the minimum and maximum values that are not outliers. Statistical comparisons Mann-Whitney test (B-E) *p<0.0.5; **p<0.01; ***p<0.001.
